# Comparative Analysis of Biomarkers in Type 2 Diabetes Patients with and without Comorbidities: Insights into the Role of Hypertension and Cardiovascular Disease

**DOI:** 10.1101/2024.01.25.577186

**Authors:** Symeon Savvopoulos, Haralampos Hatzikirou, Herbert F. Jelinek

## Abstract

**Background:** Type 2 diabetes mellitus (T2DM) are 90% of diabetes cases, and its prevalence and incidence, including comorbidities, are rising worldwide. Clinically, diabetes and associated comorbidities are identified by biochemical and physical characteristics including glycaemia, glycated hemoglobin (HbA1c), and tests for cardiovascular, eye and kidney disease.

**Objectives:** Diabetes may have a common etiology based on inflammation and oxidative stress that may provide additional information about disease progression and treatment options. Thus, identifying high-risk individuals can delay or prevent diabetes and its complications.

**Design:** In patients with or without hypertension and cardiovascular disease, as part of progression from no diabetes to T2DM, this research studied the changes in biomarkers between control and prediabetes, prediabetes to T2DM, and control to T2DM, and classified patients based on first-attendance data. Control patients and patients with hypertension, cardiovascular, and with both hypertension and cardiovascular diseases are 156, 148, 61, and 216, respectively.

**Methods:** Linear discriminant analysis is used for classification method and feature importance, This study examined the relationship between Humanin and mitochondrial protein (MOTSc), mitochondrial peptides associated with oxidative stress, diabetes progression, and associated complications.

**Results:** MOTSc, reduced glutathione and glutathione disulfide ratio (GSH/GSSG), interleukin-1β (IL-1β), and 8-isoprostane were significant (p<0.05) for the transition from prediabetes to T2DM, highlighting the importance of mitochondrial involvement. Complement component 5a (C5a) is a biomarker associated with disease progression and comorbidities, with GSH/GSSG, monocyte chemoattractant protein-1 (MCP-1), and 8-isoprostane being the most important biomarkers.

**Conclusions:** Comorbidities affect the hypothesized biomarkers as diabetes progresses. Mitochondrial oxidative stress indicators, coagulation, and inflammatory markers help assess diabetes disease development and provide appropriate medications. Future studies will examine longitudinal biomarker evolution.

## 1. Introduction

Diabetes mellitus is a chronic condition that is characterized by an absolute insulin deficiency (Type 1 diabetes mellitus, T1DM) or relative insulin deficiency from insulin resistance and decreasing insulin output (Type 2 diabetes, T2DM)^1,2^. The presence of additional chronic conditions including cardiovascular disease and hypertension as well as mental health issues have a significant impact on the clinical presentation and treatment options of T2DM. Inflammation and oxidative stress have been implicated in diabetes disease progression. However, there is a paucity of knowledge on the relationship between diabetes comorbidities with inflammation and oxidative stress. There are novel therapies that act not only on diabetes, but also on other components of metabolic syndrome, including hypertension, obesity, and hepatic steatosis. Innovative diabetes therapies like exenatide long-acting release and liraglutide are enhancing glycemic control and targeting metabolic syndrome factors like obesity and lipid abnormalities^3–5^. Liraglutide improves pancreatic beta-cell function and lipid profile, potentially mitigating atherosclerosis and cardiovascular risk^5,6^. Glucagon-Like Peptide-1 Receptor Agonists (GLP-1 Ras) optimize glucose control and may prevent cardiovascular events, promoting holistic cardio-metabolic health in diabetes care^6,7^. The development of cardiovascular disease (CVD) is significantly influenced by both diabetes mellitus (DM) and hypertension (HT)^8–12^. Epidemiological studies have estimated that the presence of T2DM leads to an approximately 2.3 times higher risk of developing CVD compared to controls with a one-third to two-thirds higher mortality associated with CVD compared to T2DM patients without CVD. Thus a more encompassing biomarker protocol for assessment of the complex heterogeneity of T2DM in the presence of comorbidities in addition to screening glucose or glycated hemoglobin (HbA1c) is required for clinical practice and identifying effective treatment options^1,2,13^.

### 1.1. Traditional biomarkers for diabetes progression and comorbidities

Traditional biomarkers for diabetes and associated complications include screening glucose, glycated hemoglobin (HbA1c), low-density lipoprotein (LDL), and total cholesterol (TC). In an oral glucose tolerance test (OGTT), screening glucose is used to define diabetes and pre-diabetes. More recently HbA1c as a clinical test has found popularity, although research has shown inconsistencies^14–16^. Similarly, in T2DM, the low-density lipoprotein (LDL) cholesterol levels vary but often are at similar levels to non-diabetes and hence are not ideal markers for diabetes with or without complications. The most typical LDL cholesterol level in diabetes is “borderline high” (130–159 mg/dl), which may increase if not addressed^17,18^. LDL cholesterol, however, is still a factor in cardiovascular risk for those with T2DM but LDL cholesterol readings may underestimate the cardiovascular risk associated with diabetes^18–20^. As for triglycerides (TG), even after adjusting for body mass index (BMI) and all other common risk factors, elevated serum TG levels are a risk factor for T2DM incidence^21^. From a clinical standpoint, elevated TG levels may be seen up to 10 years prior to T2DM diagnosis^22^. High-density lipoprotein (HDL) has also been identified as an important biomarker in diabetes progression and pathology as epidemiological research has shown that low HDL cholesterol levels are consistently linked to a higher risk of type 2 diabetes^23,24^. Extending the possible clinical biomarkers to assess diabetes progression may provide a more comprehensive assessment.

### 1.2. Oxidative stress markers for diabetes progression and comorbidities

Increased levels of reactive oxygen species (ROS), lipid peroxidation, protein carbonylation, the synthesis of nitro-tyrosine, and DNA damage are signs of increased oxidative stress in T2DM^25–27^. Hydrogen peroxide, hypochlorous acid, singlet oxygen, and radical oxygen species including superoxide anion and hydroxyl radicals can damage oxidatively crucial biological macromolecules^28^. These oxidants target unsaturated fatty acid double bonds, and lipid peroxides are produced^29^. Apolipoproteins and other plasma proteins can also be oxidized by oxygen radicals, these byproducts are more persistent than lipid peroxides for the study of diabetes progression^30^. In diabetes increased levels of inflammation, hypercholesterolemia and increased levels of LDL-cholesterol are observed that when associated with free radicals may lead to changes oxidative metabolism and CVD^31^. The cardiovascular system is one of the first systems affected by the dysglycemia in the prediabetes stage^32,33^. Increased free radicals from glucose metabolism disrupt autonomic nervous system modulation of the heart leading to cardiac autonomic neuropathy^34^. Free radicals also play an important part in lipid metabolism, inflammation and oxidative stress^35,36^.

The adaptor protein p66Shc, reduced glutathione (GSH), glutathione disulfide (GSSG), the GSH/GSSG ratio, 8-hydroxy-2-deoxyguanosine (8-OHdG) and 8-isoprostane (8-iso-PGF2α) are oxidative stress markers. Diabetic patients with high baseline levels of p66Shc have been shown to have a greater than 3-fold higher risk of diabetes associated complications during a five year follow-up compared to patients with lower p66Shc expression levels, particularly macroangiopathy^37^. However, p66shc has also been shown to decrease in prediabetes when combined in a regression model with 8-OHdG and monocyte chemo-attractant protein-1 (MCP-1), with a predictive accuracy of 89.5%^16^. But no relation between p66Shc and cardiovascular outcomes have yet been reported^37,38^. This suggests that the activity of p66shc changes during disease progression, which may be linked to the up or down regulation of other inflammatory and oxidative stress biomarkers during disease progression. 8-iso-PGF2α is considered a reliable biomarker of oxidant stress in humans^39^ and increases in the plasma of diabetic patients. A positive correlation between 8-iso-PGF2α concentration and the risk of CVD has been observed^39,40^. In addition levels of 8-iso-PGF2α are correlated with TC^41^. GSH levels decrease in patients with T2DM, with GSSG levels increasing due to the reduced GSH being oxidized^42^. Reduced synthesis and greater irreversible reactions by non-glycemic processes is a possible mechanism for the observed decrease in GSH levels^42,43^. However GSH levels vary depending on diabetes progression, presence of comorbidities and presence of other inflammatory and oxidative stress markers^16^. The findings of these previous studies imply that the de novo synthesis of GSH has been affected by CVD or HT in the presence of T2DM. Oxidative stress rises significantly in HT associated with a decrease in GSH and an increase in 8-iso-PGF2α especially when T2DM is present^41^. Finally, 8-OhdG, a DNA base alteration formed by the oxidation of deoxyguanosine, is a marker for oxidative stress in endothelial cells^44^. Compared to healthy patients, patients with prediabetes and T2DM as well as T2DM with HT have been shown to have higher urine 8-OHdG levels, which is thought to be a valuable diagnostic marker for the early diagnosis of micro-and macrovascular complications in T2DM^45,46^.

### 1.3. Mitochondrial peptide markers for diabetes progression and comorbidities

Increased screening glucose and impaired glucose metabolism also directly affects oxidative stress in mitochondria. Humanin (HN), a mitochondrial synthesized oxidative stress marker has been shown to be decreased in prediabetes and T2DM compared to control consistent with an adaptive cellular response by HN to a slight increase in screening glucose as well as being inversely linked to HbA1c^47–50^. Humanin increases insulin sensitivity, improves the survival of pancreatic beta cells, and delays the onset of diabetes^1,50^. Another mitochondrial protein, MOTSc, has also been shown to increase beta-oxidation and insulin sensitivity while directly controlling nuclear gene expression of mitochondrial biogenesis, dynamics, and function, after nuclear translocation^49,51–53^.

Other biomarkers for prediabetes, diabetes, and associated complications including inflammatory markers are C-Reactive Protein (CRP), Interleukin 6 (IL-6), Insulin-Like Growth Factor 1 (IGF-1), and Monocyte Chemoattractant Protein-1 (MCP-1)^41^. In patients with T2DM, CRP, a sensitive measure of systemic inflammation, is elevated. In addition CRP is also elevated in CVD^54^. IL-6 is a proinflammatory cytokine that, by regulating cell differentiation, migration, proliferation, and apoptosis, generates inflammation and subtly promotes the development of insulin resistance and pathogenesis of T2DM^55^. IGF-1 is a multifunctional growth factor with 50% of the same amino acids as insulin. However, insulin resistance and abnormalities in beta cell insulin production are brought on by changes in circulating IGF-1 levels^56^. Interleukin-10 (IL-10) is an immune cell-produced anti-inflammatory cytokine distinguished by its capacity to prevent macrophage activation^57^. Inflammation is caused by IL-10 or IL-10 Receptor (IL-10R) signaling defects, which may be a valid link in the development of T2DM^58^. In people with T2DM, interleukin-1beta (IL-1β) is a significant inflammatory agent^59^. Diabetes complications such as diabetic retinopathy, diabetic nephropathy, and diabetic neuropathy lead to a rise in another chemokine, MCP-1 levels in T2DM^60^. IGF-1 and MCP-1 are also involved with the pathogenesis of HT. Both a decline in IGF-1 and a rise in MCP-1 are biomarkers for inflammatory associated increases in HT patients^41^. Coagulation and fibrinolytic changes may also be associated with diabetes and diabetes progression. Complement component 5a (C5a) is an inflammatory marker and increases in diabetic patients^41^. However, plasma D-dimer levels have not been discussed extensively but higher levels may be a positive sign of diabetic peripheral neuropathy (DPN) in T2DM patients and correlate with disease progression in pre-diabetes to cardiovascular complications associated with T2DM^61,62^. Typical ranges of the afore mentioned biomarkers from the literature in control (NC), prediabetes (PreDM), and T2DM (T2DM) groups are shown in Table 1.

**Table 1.** Changes in several biomarkers types between normal (Control), prediabetes (PreDM), and type 2 (T2DM) groups based on literature.

| Biomarker | Sample | Type | Control | preDM | T2DM | References |
| --- | --- | --- | --- | --- | --- | --- |
| **IL-6 (pg/mL)** | Blood | Inflammatory | 58.56 ± 12.66 | 60.92 ± 12.82 | 60.39 ± 10.28 | <sup>63</sup> |
| **IL-1β (pg/mL)** | Blood | Inflammatory | 139.22 ± 23.23 | - | 153.26 ± 8.20 | <sup>59</sup> |
| **MCP-1(pg/mL)** | Blood | Inflammatory | 39.98 ± 15.00 | - | 65.30 ± 22.00 | <sup>64</sup> |
| **IGF-1 (pg/mL)** | Blood | Inflammatory | 199.57 ± 40.03 | 124.79 ± 27.39 | 102.03 ± 22.70 | <sup>56</sup> |
| **IL-10 (pg/mL)** | Blood | Anti-inflammatory | 229.23 ± 46.54 | 244.03 ± 53.34 | 265.04 ± 40.42 | <sup>63</sup> |
| **8-OHdG (pg/mL)** | Blood | Oxidative Stress | 177.80 ± 91.10 | 516.50 ± 260.00 | 1926.90 ± 1197.00 | <sup>45</sup> |
| **8-isoprostane (mg/mmol)** | Urine | Oxidative Stress | 0.21 ± 0.16 | 0.23 ± 0.19 | - | <sup>65</sup> |
| **GSH (mmol/L)** | Blood | Oxidative Stress | 0.85 ± 0.20 | - | 0.50 ± 0.15 | <sup>42</sup> |
| **p66Shc (pg/mL)** | Urine | Oxidative Stress | 52.30 ± 2.90 | 42.30 ± 2.30 | - | <sup>16</sup> |
| **Humanin (pg/mL)** | Blood | Mitochondrial | 1292.80 ± 56.40 | 783.70 ± 620.20 | 565.30 ± 460.50 | <sup>49</sup> |
| **MOTSc (pg/mL)** | Blood | Mitochondrial | 235.30 ± 181.60 | 154.30 ± 66.30 | 186.30 ± 130.00 | <sup>49</sup> |

**Abbreviations:** IL-6 – interleukin-6, IL-10 – interleukin-10, IL-1β – interleukin-1β , MCP-1 – monocyte chemo-attractant protein-1, IGF-1 – insulin-like growth factor 1, 8-OHdG – 8-hydroxy-2-deoxyguanosine, GSH – reduced glutathione, MOTSc – mitochondrial protein, p66Shc – adaptor protein

### 1.4. Aim of this study

In the context of T2DM, it is acknowledged that there exists a plethora of potential biomarkers for investigation^66^. This study focuses on biomarkers like HbA1c, C5a, D-Dimer, IL-6, Triglyceride, TC, HDL, LDL, 8-isoprostane, 8-OHdG, GSH/GSSG, IL-1β, IL-10, MCP-1, IGF-1, Humanin, MOTSc, and p66Shc. This emphasis is due to the study’s focus on these biomarkers and diabetes progression, particularly in the context of comorbidities. The study also uses established oxidative stress and inflammation biomarkers^41,45,50,65^. Additionally, the study includes age, cholesterol profile, and clinical characteristics. These additional variables controlled for covariates and helped identify associations. This comprehensive approach allows for a thorough analysis of the many factors that shape T2DM pathophysiology.

## 2. Materials and methods

### 2.1. Recruitment of participants

Data from 581 people attending the Diabetes Health Screening Clinic (DiabHealth) at Charles Sturt University, Albury, Australia, were chosen for analysis of blood and urine samples. This study received ethical approval from the University Human Ethics Committee, with Protocol Number 2006-042, and all participating patients provided written informed consent, which included consent for publication of results. The preparation of this manuscript adhered to the STROBE (Strengthening the Reporting of Observational Studies in Epidemiology) Guidelines, ensuring comprehensive and standardized reporting of our observational cross-sectional study^67^. There were no requirements for exclusion from this study as patients attended a diabetes complications progression research initiative clinic and hence any clinical data relevant to patient health was collected. All participants had their age, gender, HbA1c, C5a, D-DIMER, IL-6, Trigs, TC, HDL cholesterol, LDL cholesterol, 8-isoprostane, 8-OHdG, GSH, GSSG, GSH:GSSG ratio, IL-1β, IL-10, MCP-1, IGF-1, Humanin, MOTSc, and p66Shc recorded. A control group with a screening glucose of < 5.6 mmol/L, a prediabetes group with a screening glucose of 5.6–6.9 mmol/L, and a type 2 diabetes group with screening glucose ≥ 7 mm/L were compared. The study’s nature was to focus on oxidative stress biomarkers and diabetes progression, particularly in the context of comorbidities, over a 10 year period, utilizing data only from the patients’ first admission. To determine the sample size, an ANOVA power analysis was conducted, with three groups representing ‘no comorbidities,’ ‘preDM,’ and ‘T2DM.’ We assumed an effect size of f=0.5, a significance threshold of α=0.05, and a desired statistical power of 0.8. As a result, the calculated sample size for each group was determined to be 13.

#### Apparatus

Accu-Chek® (Roche Australia Pty Ltd.) measured screening blood glucose. Blood preparation centrifugation was done with a UNIVERSAL 32R (Hettich Zentrifugen). The photometric analysis of biomarkers in urine was carried out with the assistance of a Thermo Scientific Multiskan FC (Fisher, China).

### 2.2. Sample preparation

Midflow urine samples were collected to measure all oxidative stress and inflammatory markers. Screening glucose, HbA1c and cholesterol profile was determined by the local pathology laboratory.

### 2.3. Measurement of oxidative stress

The Glutathione EIA Kit (Cayman Chemical, USA) measured erythrocyte GSH and GSSG using glutathione reductase. The formation of yellow 5-thio-2-nitrobenzoic acid (TNB), which is directly proportional to the total GSH concentration in the sample, is caused by the reaction of 5,5′-dithio-bis-2-nitrobenzoic acid, also known as DTNB. Before the experiment, samples and standards were treated with 1% of 1 M 2-vinylpyridine for 60 min at room temperature to derivatize free GSH and detect GSSG exclusively. GSH and GGSG concentrations and deproteination were calculated using the End Point Method. Northwest (USA)’s urinary isoprostane ELISA kit uses a competitive ELISA strategy to measure the amount of 8-isoprostane in samples and standards. After adding the horseradish peroxidase substrate, 8-isoprostane in samples and standards inversely correlates with the blue color development, which turns yellow after acid termination. 450 nm is absorbance.The Human SHC-Transforming Protein 1 ELISA kit (CUSA-BIO; Flarebio Biotech LLC) was used to measure p66shc because, according to the kit’s creators, it primarily detects Shc1 and its isoform p66shc. A commercially available extraction-free EIA kit was used to measure MOTS-c concentrations in accordance with the manufacturer’s suggested procedure. As for Humanin, the Elisa analysis (Lot No. K11064644) was utilized from Elisakit.com (Adelaide, Australia).

### 2.4. Measurement of endothelial dysfunction

An 8-hydroxy-2-deoxy guanosine EIA Kit (Cayman Chemical, USA) measured urine 8-OHdG. An anti-mouse IgG-coated plate and an 8-OHdG-acetylcholinesterase conjugate tracer increased sensitivity and decreased variability in this competitive test. 8-OHdG-enzyme conjugate (tracer) and sample 8-OHdG compete for monoclonal antibody. After the tracer-antibody complex binds to the pre-coated anti-mouse IgG, AChE-substrate (acetylcholine and 5,5′-dithio-bis-2-nitrobenzoic acid) is added to produce a yellow 5-thio-2-nitrobenzoic acid that can be detected spectrophotometrically at 412 nm and is inversely proportional to the amount of 8-OHdG in the sample. To standardize 8-isoprostane and 8-OHdG results, Albury’s Dorevitch Pathology Laboratory quantified urine creatinine. The automated AxSYM® system (Abbott Laboratories, USA) measured plasma homocysteine with a two-step reaction: 1. dithiothreitol reduction of protein-bound homocysteine, and 2. enzymatic conversion of free homocysteine to S-adenosylhomocysteine with adenosine. Fluorescence polarization immunoassay detects reaction 2 product S-adenosylhomocysteine.

### 2.5. Measurement of inflammation

Plasma IL-6 was tested using a double-antibody sandwich ELISA Interleukin-6 (human) EIA Kit that was manufactured by Cayman Chemical in the United States. An acetylcholinesterase:IL-6 Fab’ combination binds to a particular epitope on the IL-6 molecule, whereas a pre-coated monoclonal antibody binds to free IL-6 in samples. Both of these bindings are carried out by the IL-6 molecule. The AChE-substrate, which is composed of acetylcholine and 5,5′-dithio-bis-2-nitrobenzoic acid, is responsible for initiating the enzymatic synthesis of the yellow 5-thio-2-nitrobenzoic acid, which has an absorption peak at 412 nm. IL-6 levels determine color development. The serum CRP concentrations were obtained by the Dorevitch Pathology Laboratory in Albury, New South Wales.

### 2.6. Measurement coagulation and fibrinolysis

The Human C5a ELISA Kit II (BD Biosciences, USA) measured plasma C5a using a sandwich ELISA test using a human C5a-specific monoclonal antibody pre-coated on microplates. The second antibody that binds to immobilized C5a is streptavidin horseradish peroxidase conjugate, which reacts with TMB to create blue. Phosphoric acid inhibits the process and makes it yellow. Absorbance at 450 nm determines C5a content in the first sample. The Albury-based Dorevitch Pathology Laboratory provided D-Dimer readings.

### 2.7. Statistical analysis

Microsoft Excel (Office 365, Microsoft) was used to analyze descriptive data and is presented as mean ± standard deviation (x ± SD). Statistical analysis was carried out with Python with the Pingouin, which is an open-source statistical package^68^. A Kruskal-Wallis test was conducted to evaluate whether there were any notable variations in biomarker levels between the groups^69^. Additionally, a Mann Whitney post hoc test was utilized to compare differences between any 2 groups^70^. Significant data was defined as a p-value < 0.05. In this study, linear discriminant analysis (LDA) was employed to identify the most significant biomarkers and predict the outcomes. Data from a D dimensional feature space was projected using LDA into a D’ (D>D’) dimensional space in order to increase variability across classes while minimizing variability within classes^71^. Standard scaling was a crucial preprocessing step in LDA to guarantee robust, interpretable, and unaffected analysis based on the input feature scale. In terms of feature importance, we assessed it by considering the features with higher absolute coefficient values in the LDA model. Important features were compared with the p-values of Kruskal-Wallis and Mann Whitney tests. LDA classifications of various populations using varying degrees of k-fold cross-validation were used. Accuracy estimations were made of the LDA area under the curve (AUC) of the receiver operating characteristic (ROC) curves between the two classes without and with HbA1c, as well as the accuracies between the three classes without HbA1c, and with HbA1c.

## 3. Results

This section is divided into four parts. The first part is focused on the healthy control group compared to prediabetes and diabetes without comorbidities, the second part includes patients with HT and CVD, the third is devoted to patients with HT only, and the fourth part shows results of patients with CVD only. Each part shows the main significant biomarkers of the three groups; control/no diabetes, prediabetes (preDM), and type 2 diabetes (T2DM). Kruskal Wallis and Mann Whitney analyses were used to estimate the difference between multiple groups and between two groups, respectively (the expanded Tables with the results of the analyses are in the Appendix). In each part, by using LDA, the most critical biomarkers are shown for two groups (at least absolute normalized eigenvector 0.1). Area under the curve (AUC) of receiver operating characteristic (ROC) curves are provided, and the accuracies for all classes. The detailed ROC curves are included in the Appendix. In feature importance and LDA, the classes studied were: A.1.) Control/no diabetes – preDM without HbA1c, A.2.) Control/no diabetes – preDM with HbA1c, B.1.) preDM – T2DM without HbA1c, B.2.) preDM – T2DM with HbA1c, B.1.) Control/no diabetes – T2DM without HbA1c, C.2.) Control/no diabetes – T2DM with HbA1c, and D.1.) Control/no diabetes -preDM – T2DM without HbA1c, D.2.) Control/no diabetes - preDM – T2DM with HbA1c.

### 3.1. Important features in the healthy population

Table A1 in the Appendix summarizes the clinical variables measured in the different healthy groups. Significant differences between the three groups (using the Kruskal-Wallis test) were observed for HbA1c, D-Dimer, Triglycerides, HDL, GSSG, IL-1β, IL-10, IGF-1, Humanin, MOTSc, and p66Shc. MCP-1, and 8-isoprostane showed differences between control and T2DM groups, but they were not significant based on the Kruskal-Wallis analysis. Comparing the three groups further it can be observed that IL-10, IGF-1, and Humanin were highest in the control group. Patients with preDM had the highest HDL, IL-1β, and MOTSc levels. Lastly, HbA1c, D-Dimer, Triglycerides, and p66Shc levels were greater in T2DM group compared to control and preDM. Before proceeding to the classification, biomarker importance with respect to two and the three groups without HbA1c (A) or with HbA1c (B) are included in Figure 1.

**Figure 1:**
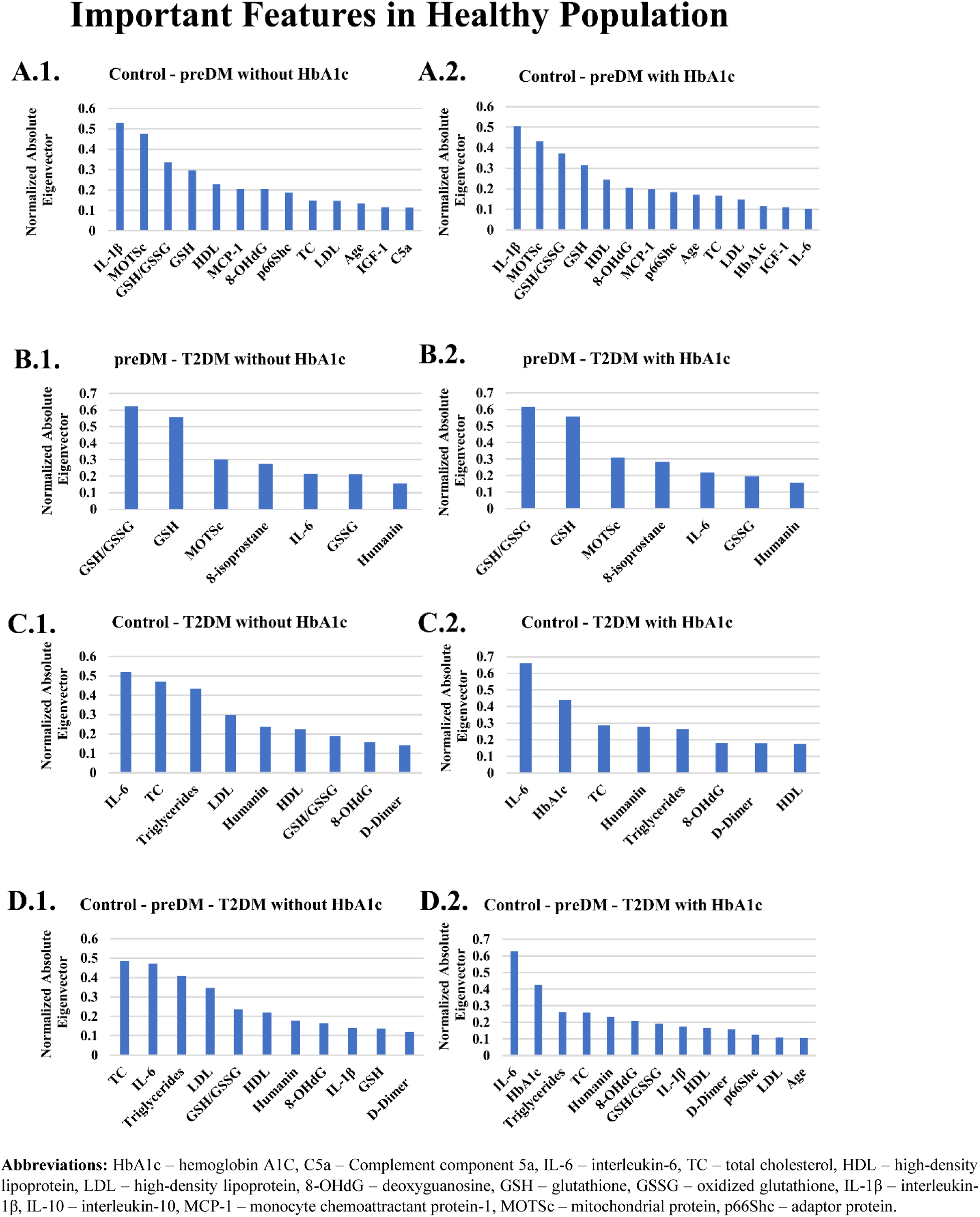
Important features in the healthy population with the use of LDA

The most dominant biomarkers with respect to absolute normalized eigenvector ≥ 0.1, were also the most significant as shown in Figures 1. A.1.-D.2.. The only difference is IL-6, which is the most dominant biomarker for the control group with T2DM (Figure 1. C.1.-C.2.), and with preDM and T2DM (Figure 1. D.1.-D.2.).

LDA classifications with different k-fold cross-validation are plotted in the ROC curves in Figure A1 in the Appendix. The AUC of the ROC curves between two classes without and with HbA1c were above 0.98 for 5-fold cross validation, and the accuracies between the three classes without HbA1c, and with HbA1c are both equal to 0.94±0.02. Thus, classification can be achieved successfully between two groups, and three groups as the accuracies were high with or without HbA1c.

### 3.2. Important features in the population with HT and CVD

Table A2 in the Appendix lists the clinical parameters that were evaluated for the three groups in with HT and CVD. The demographic factors for HbA1c, C5a, D-Dimer, Triglycerides, TC, HDL, LDL, 8-isoprostane, GSSG, GSH/GSSG, IL-10, MCP-1, Humanin, and MOTSc showed significant differences according to the Kruskal-Wallis and Mann Whitney tests. The levels of D-Dimer, HDL, MCP-1 of the no diabetes group compared to the other two groups were higher. PreDM patients had higher levels of LDL, and GSH/GSSG compared to no diabetes and T2DM groups. Finally, the levels of HbA1c, C5a, Triglycerides, 8-isoprostane, GSSG, IL-10, Humanin, and MOTSc were higher in T2DM patients compared to no diabetes and preDM groups. HbA1c, and triglycerides were elevated in T2DM patients of both the healthy group (Table A1) and the group with HT and CVD (Table A2).

Figure 2 A.1.-D.2. depict the importance of biomarkers divided into two or three groups without HbA1c (A) or with (B) HbA1c before classifying the population of both HT and CVD. The most prevalenbiomarkers (p<0.05 in Table A2 in the Appendix) were the most significant using Kruskal-Wallis and Mann-Whitney analyses. Only GSH, and IGF-1 were the two additional features which showed an importance in the HT and CVD population independent from Kruskal Wallis and Mann Whitney analyses. The decreased GSH, emerged as a significant biomarker when the preDM group was compared to the T2DM group (Figure 2 B.1. and B. 2.). In addition, when the preDM group was compared to individuals who had T2DM, IGF-1 declined, and emerged as a relevant biomarker (Figure 2 B.1. and B.2.).

LDA classifications of the HT and CVD population with different k-fold cross-validation were plotted in Figure A2 of the Appendix. The LDA achieved AUC of the ROC curves between two classes without and with HbA1c were above 0.83 for 5-fold cross validation, and the accuracies between the three classes without HbA1c, and with HbA1c are both equal to 0.85±0.03.

**Figure 2:**
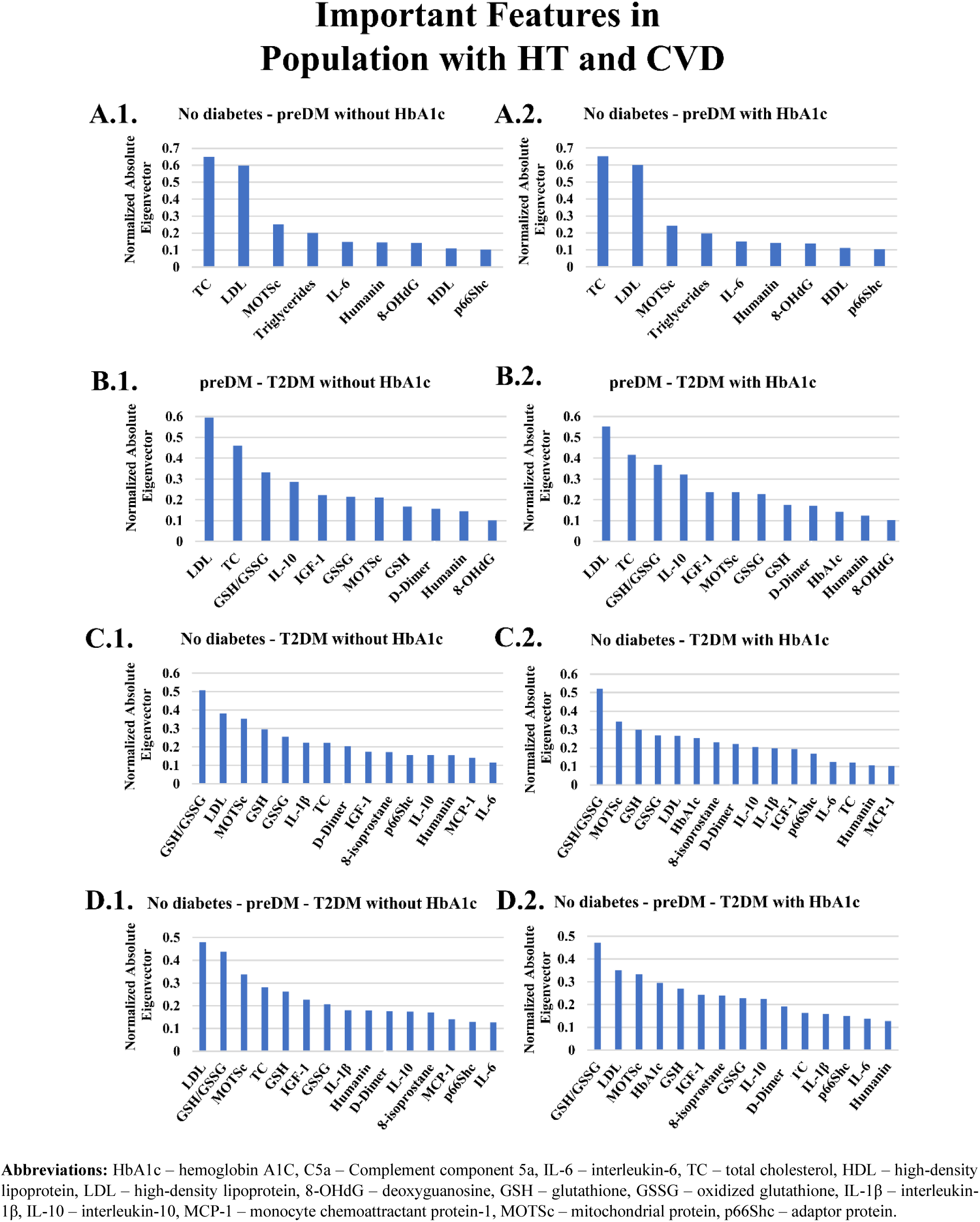
Important features in the HT and CVD population with the use of LDA.

### 3.3. Important features in the population with HT only

Table A3 in the Appendix lists the clinical parameters that were evaluated in the various groups in the population that had only HT. The demographic factors for HbA1c, C5a, Triglycerides, TC, LDL, 8-isoprostane, 8-OHdG, GSH, GSSG, GSH/GSSG, IL-1β, IL-10, MCP-1, IGF-1, Humanin, and MOTSc showed significant differences according to the Kruskal-Wallis and Mann Whitney tests (p<0.05). The no diabetes group showed an increased TC, LDL, GSH, GSSG, IL-1β, IL-10, and MCP-1 compared to the other groups. Patients with preDM showed increased 8-isoprostane, 8-OHdG, and IGF-1 values compared to the no diabetes and T2DM groups. Finally, T2DM patients had increased levels of HbA1c, C5a, Triglyceride, GSH/GSSG, Humanin, and MOTSc. Triglycerides and HbA1c remained elevated, similar to those of the T2DM patients in the healthy group in Table A1 and the group with HT and CVD in Table A2.

Figure 3. A.1-D.2. depict the importance of biomarkers for two and the three groups without (A) or with (B) HbA1c before classifying the population with both HT and CVD. The most prominent biomarkers were the most different in both Kruskal-Wallis and Mann-Whitney analyses (Table A3 in the Appendix). The only difference is in IL-6, and HDL which were the most dominant biomarkers due to their lower and maximum values, respectively for the comparison of the no diabetes group with the other groups. LDA classifications with different k-fold cross-validation are plotted in Figure A3. The LDA successfully classified the groups as this method achieved AUC of the ROC curves between two classes without and with HbA1c above 0.83 for 5-fold cross validation, and the accuracies between the three classes without HbA1c, and with HbA1c were both equal to 0.87±0.04.

**Figure 3:**
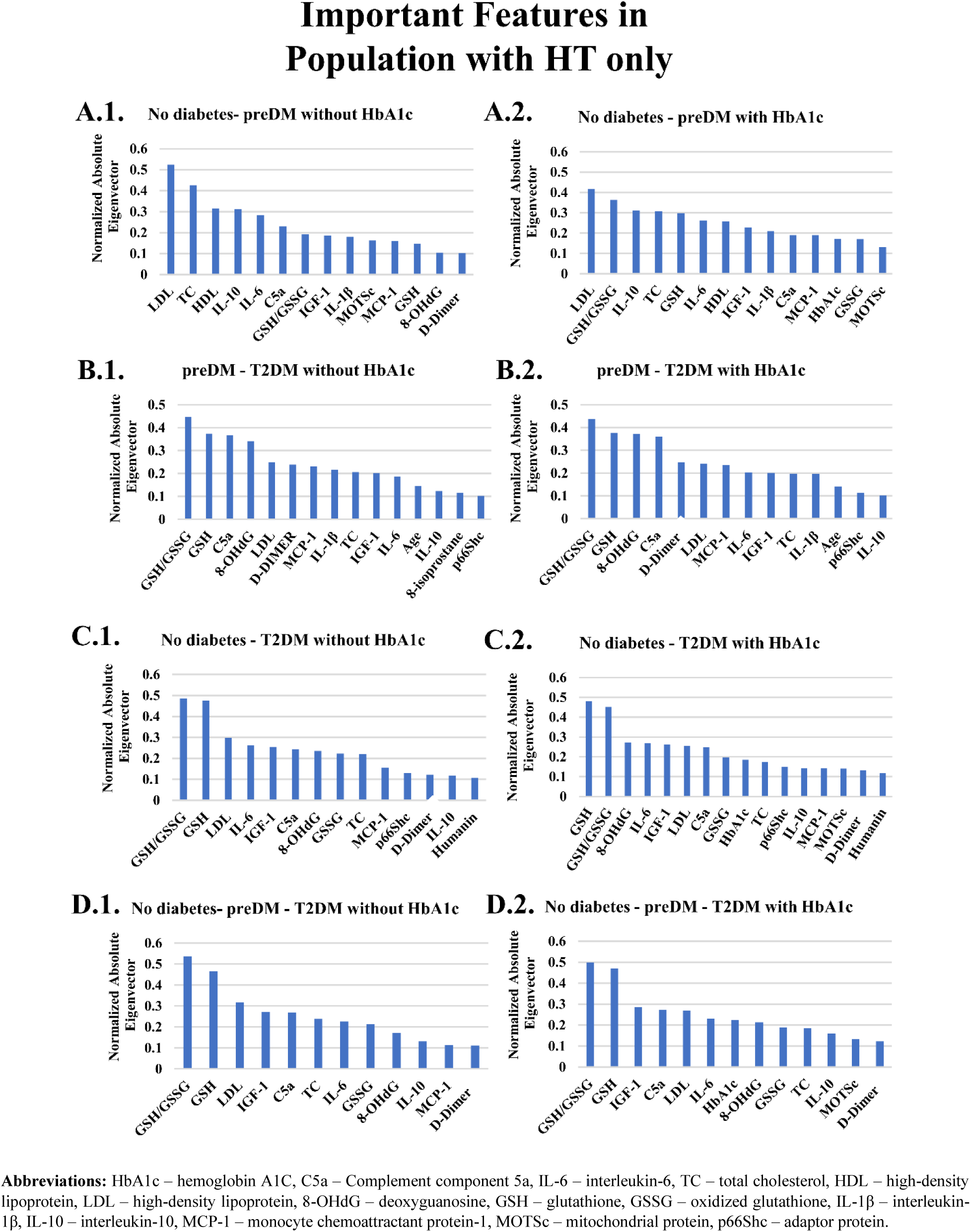
Important features in the HT population with the use of LDA.

### 3.4. Important features in the population with CVD only

The clinical features evaluated in the various CVD groups are summarized in Table A4 in the Appendix. The Kruskal-Wallis test identified significant differences in the demographic factors for HbA1c, IL-6, TC, HDL, LDL, 8-OHdG, GSH, GSSG and IL-10. The levels of TC, HDL, and LDL were significantly higher in the no diabetes group compared to preDM and T2DM patients. In contrast, preDM patients had higher levels of IL-6 compared to the no diabetes and T2DM groups (p<0.05). Finally, the levels of HbA1c, GSH, and GSSG were higher in T2DM patients (p<0.05). These differences in levels of biomarkers substantiate a dynamic, fluid representation of the disease progression.

Figure 4.A.1-D.2. illustrate the importance of biomarkers for no diabetes, preDM and T2DM without HbA1c (A) or with (B) HbA1c. Only MCP-1 (Figure 4 A.1.-A.2.), 8-isoprostane (Figure 4 B-D.1.-2.) and GSH/GSSG (Figure 4 A.1.-2, C.1.-2, and D.1.-2) show a discernible difference and increase from no diabetes to T2DM.

Figure A4 displays the LDA classifications with various k-fold cross-validations. The LDA achieved AUC of the ROC curves between two classes without and with HbA1c were above 0.83 for 5-fold cross validation, and the accuracies between the three classes without HbA1c, and with HbA1c are both equal to 0.85±0.09. Because the accuracy rates were high with or without HbA1c, classification between the two groups and three groups can be conducted accurately.

**Figure 4:**
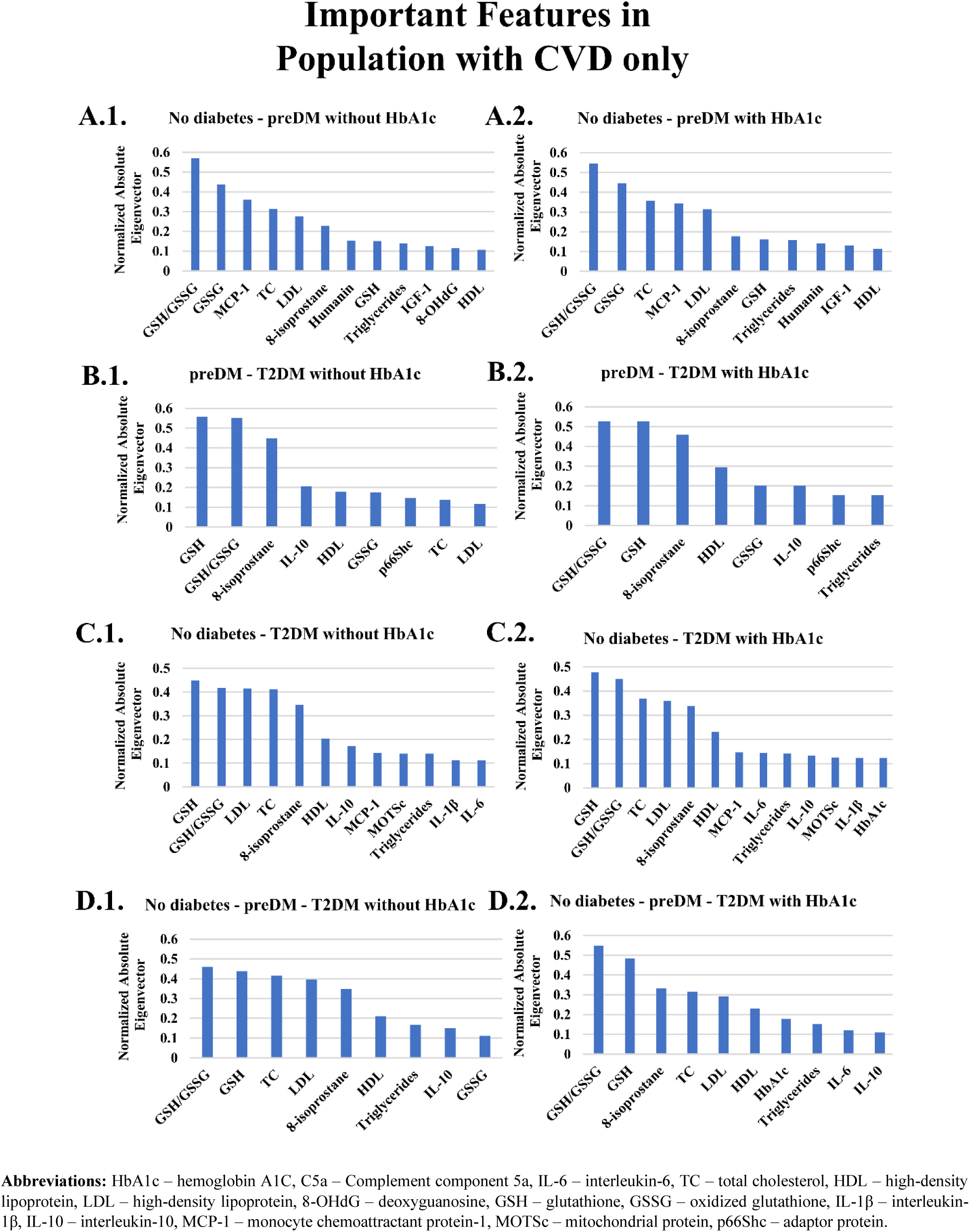
Important features in the CVD population with the use of LDA.

## 4. Discussion

The overall objective of this research was to study the changes in a set of measurable biomarkers between control and prediabetes, prediabetes to T2DM and control to T2DM and to classify patients based on their first-attendance data at the clinic. Diverging from previous analysis, this study presents the importance of biomarkers at specific stages of T2DM progression and allows visualization of possible changes in the metabolic pathways and interactions. More precisely, our analysis (1) uses biomarker measurements from the first attendance of the patient, (2) considers the importance of the biomarkers at each stage of the disease progression, (3) considers patients with CVD, and HT comorbidity, and (4) predicts the clinical T2DM stage outcome with high accuracy. Our study aligns with the ongoing efforts for improving the diagnosis of T2DM, assessing advanced prediction schemes and finally, for realistic and optimized disease treatment, which is the next step in this research area.

Our results suggest in agreement with others that HbA1c may not be a good marker for identifying diabetes progression if comorbidities including CVD and/or HT are present. HbA1c when included in the analysis tended to play a minor role in terms of contributing to the model. However, the consistently greater increase of HbA1c in the CVD/HT comorbidity highlights its association with increased risk of CVD morbidity and mortality^72,73^.

Our results revealed some novel and clinically useful associations for diabetes progression in the presence or absence of CVD and HT as comorbidities. Several of the significant markers were expected including triglycerides, HDL, IL-1β, IL-10 and IGF-1. D-Dimer has not been investigated extensively. However, previous work of ours has shown a relationship between diabetes progression in the presence of macrovascular disease and more recently D-dimer has been shown to be a promising marker for peripheral neuropathy^61,62^. GSSG is a sensitive oxidative stress marker, which reflects possible decreasing synthesis of GSH and increased irreversible utilization by reactive oxygen species reactions^42^.

Of interest are the results for Humanin and MOTSc, which are mitochondrial peptides associated with oxidative stress that have not been investigated together with reference to diabetes progression and associated complications^74^. In agreement with Ramanjaneya, we observed changes in both Humanin and MOTSc with diabetes progression and correlations with HbA1c and HDL^49^. In addition, p66Shc, which is coded for in the nucleus but also associated with oxidative stress by modulating the mitochondrial electron transport chain^16,75^. However, for the CVD/HT patients C5a, TC, LDL, 8-isoprostane, GSH/GSSG, and MCP-1 became important indicating significant changes in the pathophysiology and associated biomarkers with disease progression and the influence of CVD and HT. As for the patients with HT only, the importance of C5a, 8-OHdG, GSH/GSSG, IGF-1, became significant in the model in addition to the well documented role of cholesterol. Gouaref et al.^76^ reported similar results between a control group and patients with T2DM, hypertension, and hypertension. In contrast patients with CVD differed significantly only in TC, HDL, LDL, 8-OHdG, GSH, GSSG, and IL-10 indicating the much more prominent role of cholesterol in cardiovascular disease.

### 4.1. Diabetes progression without comorbidity

The dynamics of the biomarkers at different transitions in the group with no comorbidities indicated a strong association with oxidative stress markers rather than cholesterol or inflammatory markers. MOTSc, and GSH/GSSG as well as IL-1β were significant for the transition from control to preDM, agreeing with previous research^77^. For the transition from preDM to T2DM GSH/GSSG, MOTSc, and 8-isoprostane were significant and highlighting the importance of mitochondrial involvement^78^. When the control group was compared to the T2DM group, IL-6 appeared in the model as an inflammatory marker^79^ in addition to. a switching from MOTSc to Humanin, which indicates their possible different roles in the pathophysiology of complex disease and may reflect the specific functions and hormone-type activity of these mitochondrial peptides^80^. MOTSc stimulates cellular glucose uptake and may be an early response to increased screening glucose at the prediabetes phase. In contrast Humanin reduces inflammatory markers, interacts with IGF-1 and has been shown to decrease with disease progression^81^.

### 4.2. Diabetes progression with comorbidity

In the group with both HT and CVD, a change in biomarkers with disease progression was again observed. For the change from no diabetes to preDM apart, from cholesterol, MOTSc was the most important marker, whereas GSH/GSSG, IL-1β and IL-10 became associated with further disease progression if HbA1c was included in the model. Otherwise, 8-isoprostane was an additional oxidative stress marker. This results also indicates the variability of HbA1c with T2DM and presence of comorbidities^82^. Kim et al. showed that MOTSc improves beta-oxidation and plasma GSSG was lower in a MOTS-c injected animal model providing further evidence for the complex interaction of biomarkers with disease progression highlighted in the current study^74,83^. Thus, at the prediabetes stage identification of a battery of risk factors including oxidative stress markers may provide an optimal opportunity for successful multifactorial intervention including life-style factors, and polypharmacy if required^84^. MOTSc inhibits inflammation and blocks cellular apoptosis amongst other functions.

When the biomarker changes in the diabetes and HT group was the coagulation marker C5a and the fibrinolytic marker, D-dimer appeared in the models Complement C5a mediates pro-inflammatory responses and is elevated in diabetes as well as playing a role in diabetic nephropathy^85^. The current study extends our understanding of the role of C5a as direct associations of this biomarker was observed with disease progression and presence of comorbidities, providing a useful timeline in terms of the biochemical milieu changes associated with disease progression. D-dimer has previously been reported to be already elevated in prediabetes and diabetic peripheral neuropathy has recently been associated with increased levels of D-dimer alongside fluctuations in HbA1c^61,86^.

The most important biomarkers in the shift from no diabetes to preDM, in the CVD comorbidity group, were GSH/GSSG, MCP-1, GSH/GSSG, and 8-isoprostane in transition from preDM to T2DM, whilst for the shift from no diabetes to T2DM GSH/GSSG, and 8-isoprostane were the most important biomarkers. Changes in MCP-1 were not significantly different either being slightly increased in the prediabetic group and then slightly decreased in the T2DM group. 8-isoprostane is a marker of lipid oxidation and onset of CVD, specifically associated with endothelial dysfunction and inflammation^87^. However, in our study the increase in 8-isoprostane was not significant. Previous work of ours reported on the importance of D-dimer and the correlation with GSH levels associated with disease progression and including CVD as comorbidity^86^. In the current study prediabetic and T2DM CVD patients had a lower GSH/GSSG ratio than healthy controls, indicating more oxidative stress associated with disease progression. Hypoglycemic and cardiovascular disease medicines’ antioxidant effects may explain the prediabetes group’s small GSH rise. Another possibility is that increased GSSG production due to oxidative stress, likely caused by erythrocyte ROS levels, stimulates GSH production^65^.

Research into the pathophysiological mechanisms related to diabetes progression with CVD has shown the important role of oxidative stress and use of medication in preventing and treating diabetes associated CVD^88^. Early intervention is warranted for patients with CVD as changes in inflammation and oxidative stress are already present at the prediabetes stage and possibly prior to any clinical identification of any CVD risk including obesity^88–90^.

In addition, previous research has shown that CVD patients without diabetes, with prediabetes or diabetes had higher levels of GSSG and 8-isoprostane^91^, and MCP-1 than healthy controls^92^ as well as increased LDL^93,94^, and TC cholesterol^60,95^.

### 4.3. Influence of medication on biomarker levels

Some of the inconsistent results with laboratory studies, stem form our research taking place in a diabetes complications screening clinic, where patients were on different types of medication that may influence levels of cholesterol and other biomarkers. Prediabetes and T2DM may have higher LDL and TC values and lower HDL levels than control individuals^96^. In our group, the drop in cholesterol profile is probably due to medication. Depending on the type of antihypertensive used, the literature has shown either beneficial effects or negative effects of antihypertensive use^97^. Kasiske et al found β-blockers with intrinsic sympathomimetic activity and cardioselectivity reduction of TC and LDL-C. α-blockers beneficially affected TC, LDL-C, TG, and in younger individuals, HDL^98^. ACE inhibitors reduced TG in diabetics, TC. Direct vasodilators reduced TC and LDL, and increased HDL. Alternatively, diuretics increase TG and TC and have also a negative impact on LDL. β-1 selective and nonselective β-blockers may increase TG and lower HDL. Compared to control group, prediabetic and T2DM patients with HT may have lower IL-10 levels, indicating inflammation^99^. T2DM and prediabetic patients with HT may have higher IL-6 levels than control group, indicating elevated inflammation in advanced disease stages^100^. Prediabetic and T2DM HT patients have been shown to have lower GSH/GSSG ratios, indicating increased oxidative stress^65^. Similarly in the current study with a larger sample and comorbidities, the prediabetic patients with HT and greater 8-OHdG levels than control and T2DM, indicate more DNA damage in prediabetes, which is reduced due to antioxidant activity of polypharmacy^101^. Atenolol and metformin are widely used antioxidant, and antidiabetic medicines, respectively^102–104^. T2DM patients with HT have lower IGF-1 levels than controls and prediabetics, suggesting impaired growth and repair mechanisms^105^. Similarly, T2DM patients with HT and CVD have higher LDL and TC levels and a lower GSH/GSSG ratio than healthy controls and pre-diabetics. Again, the decreased LDL and TC levels in our groups might be due to the medication. The prevalence of MOTS-c may protect T2DM patients with HT and CVD by improving mitochondrial activity, insulin sensitivity, oxidative stress, inflammation, and endothelial function in agreement with Mohtashami et al. (2022)^74^. Also, MOTS-c plays a role in lipid metabolism, blood pressure management, and cardiac function^106^.

Our results highlight the importance of assessing disease progression in terms of multifactorial involvement by applying linear discriminant analysis and variable importance assignment. The majority of our results reflect previous findings for studies that investigated single inflammatory or oxidative stress markers but highlight the importance of comparing of a single biomarker between stages of disease development and included in the model of disease progression.

### 4.4. Data analytics in diabetes progression

Utilizing the feature importance of T2DM, established methods have been employed by using different sets of inputs^107,108^. In the work of Bernadini et al., a multiple instance learning boosting (MIL-Boost) algorithm was used for creating a predictive model of worsening insulin resistance in T2DM in terms of Triglyceride-glucose index (TyG)^109^. The algorithm was able to extract hidden patterns from past electronic health record temporal data. Triglycerides, HDL cholesterol, and total cholesterol were among the most important aspects of the MIL-Boost experimental design, and these were included in our analysis, as well. Jelinek et al.^15^ investigated whether additional biomarkers could be used in conjunction with HbA1c to improve diagnostic accuracy in T2DM if HbA1c levels are less than or equal to the current cutoff value of 6.5%. They concluded that oxidative stress marker 8-hydroxy-2-deoxyguanosine (8-OHdG) and interleukin-6 (IL-6) improved classification accuracy. Finally, Hathaway et al. developed a model for precision medicine by applying machine-learning algorithms to predict the development of T2DM using various cardiac indicators, as T2DM plays a crucial role on cardiovascular disease^110,111^.

Our study adapts to each of the several case studies in a meaningful way, and the parameters and variables follow physiologically appropriate ranges. The proposed framework was created using a number of biomarkers and comorbidities (HT and CVD). The created algorithm, however, may enable the detection of pre-diabetes, diabetes, and related comorbidities if patient data from our study are included. One of the limitations of this research pertains to the small number of patients within certain groups. Future studies should consider augmenting the sample size for these specific patient cohorts to enhance the robustness of findings. Additionally, future research of ours will now focus on the dynamics of biomarker changes during the various stages of Type 2 diabetes mellitus progression.

## 5. Conclusions

Measures of the proposed biomarkers has shown the influence of comorbidities on biomarker involvement as a function diabetes progression. The observed changes of mitochondrial oxidative stress markers as well as coagulation and inflammatory maker changes allow an improved assessment of diabetes disease progression and decisions on appropriate individualized medication prescription. In upcoming studies, the longitudinal biomarker evolution will be assessed.

## Declarations

### Ethics approval and consent to participate

University Human Ethics Committee, Protocol Number 2006-042. All patients gave written informed consent to participate.

### Consent for publication

All participating patients provided written informed consent, which included consent for publication of results.

### Author Contribution

Conceptualization (SS, HH, HJ), Data Curation (SS, HJ), Formal Analysis (SS, HH), Investigation (all authors), Methodology (SS, HJ), Resources (all authors), Software (SS), Supervision (HJ), Validation (SS), Writing original draft (SS, HJ), Writing review and editing (all authors).

## Acknowledgments

The authors which to acknowledge the assistance of Bev DeJong in the data collection and the assistance of Charles Sturt University funding. Part of this work was undertaken by visiting students through the Deutscher Austausch Dienst.

## Availability of data and material

Please contact corresponding author for available data.

