## Appendix for "Comparative Analysis of Biomarkers in Type 2 Diabetes Patients with and without Comorbidities: Insights into the Role of Hypertension and Cardiovascular Disease"

The Appendix is divided into four parts. The first part is focused on the healthy control group compared to prediabetes and diabetes without comorbidities, the second part includes patients with HT and CVD, the third is devoted to patients with HT only, and the fourth part shows results of patients with CVD only. Each part contains a table (Tables A1-A4) that shows the main significant biomarkers of the three groups; control/no diabetes, prediabetes (preDM), and type 2 diabetes (T2DM). Kruskal Wallis and Mann Whitney analyses were used to estimate the difference between multiple groups and between two groups, respectively. As for the classification, the ROC response of various datasets produced by k-fold cross-validation is displayed in Figures A1–A4 below. The ROC curves for several classes of the healthy population without or with HbA1c are shown in Figure A1. Similar to Figure A1, Figure A2 showed the ROC curves for the HT and CVD patients. Additionally, Figure A3 contains a classification analysis of patients who solely have HT. Last but not least, Figure A4 contains the ROC analysis for just the individuals with CVD. When the training set is divided into distinct subsets, the mean area under the curve (AUC) can be calculated by altering the k-fold cross validation. This roughly illustrates how changes in the training data affect the classifier output and how distinct the splits produced by k-fold cross-validation are from one another. The categorization of our data revealed that the control/no diabetes. with preDM classes were the only two that significantly changed when the k-fold was changed. The AUC between these two classes and all classes were maximum when k=5 was used. The accuracy was applied when the three classes were used. Overall, the classification process for k=5 achieved an AUC of more than 80%, which is considered to be successful.

a) Statistical analysis and classification ROC curves for the healthy population

**Table A1:** Characteristics and biomarker results of healthy population in control, preDM, and T2DM subjects

|  |  |  |  | **Kruskal Wallis** | **Mann Whitney** | | |
| --- | --- | --- | --- | --- | --- | --- | --- |
|  | **Control** | **preDM** | **T2DM** | **All Classes** | **Control – preDM** | **Control – T2DM** | **preDM – T2DM** |
|  | **Mean ± SD** | **Mean ± SD** | **Mean ± SD** | **p-value** | **p-value** | **p-value** | **p-value** |
| **N** | 123 | 21 | 12 |  |  |  |  |
| **Age (yr)** | 55.21 ± 11.55 | 57.76 ± 9.32 | 58.92 ± 8.33 | 0.296 | 0.309 | 0.199 | 0.694 |
| **HbA1c (%)** | 5.59 ± 0.27 | 5.82 ± 0.28 | 6.89 ± 1.02 | **0.000***** | **0.001**** | **0.000***** | **0.007**** |
| **C5a (ng/mL)** | 14.73 ± 15.29 | 12.65 ± 7.99 | 15.27 ± 7.26 | 0.456 | 0.968 | 0.198 | 0.442 |
| **D-Dimer (μg/L)** | 0.38 ± 0.33 | 0.32 ± 0.08 | 0.60 ± 0.28 | **0.007**** | 0.083 | **0.008**** | **0.007**** |
| **IL-6 (pg/mL)** | 15.23 ± 7.25 | 16.50 ± 6.85 | 41.64 ± 43.72 | 0.478 | 0.433 | 0.328 | 0.536 |
| **Triglyceride (mmol/L)** | 1.12 ± 0.62 | 1.04 ± 0.36 | 3.15 ± 1.81 | **0.000***** | 0.820 | 0.420 | 0.348 |
| **TC (mmol/L)** | 5.09 ± 0.76 | 5.27 ± 0.77 | 4.82 ± 0.94 | 0.421 | 0.335 | 0.394 | 0.294 |
| **HDL (mmol/L)** | 1.39 ± 0.31 | 1.57 ± 0.31 | 1.16 ± 0.27 | **0.001**** | **0.005**** | **0.011*** | **0.001**** |
| **LDL (mmol/L)** | 3.19 ± 0.66 | 3.23 ± 0.76 | 2.99 ± 0.57 | 0.666 | 0.820 | 0.420 | 0.348 |
| **8-isoprostane (mg/mmol)** | 2.62 ± 4.46 | 1.21 ± 0.56 | 9.09 ± 8.68 | 0.149 | 0.941 | 0.047 | 0.181 |
| **8-OHdG (mg/mmol)** | 153.31 ± 95.64 | 151.45 ± 53.26 | 123.87 ± 66.43 | 0.296 | 0.413 | 0.235 | 0.101 |
| **GSH (mmol/L)** | 1649.49 ± 289.58 | 1722.77 ± 392.78 | 1703.90 ± 53.58 | 0.887 | 0.883 | 0.679 | 0.508 |
| **GSSG (mmol/L)** | 298.65 ± 73.41 | 316.32 ± 61.73 | 344.31 ± 82.06 | **0.032*** | 0.060 | **0.046*** | 0.338 |
| **GSH/GSSG (-)** | 5.79 ± 1.59 | 5.93 ± 2.63 | 5.29 ± 1.43 | 0.208 | 0.446 | 0.070 | 0.836 |
| **IL-1β (pg/mL)** | 6.76 ± 6.10 | 15.14 ± 10.61 | 11.29 ± 4.07 | **0.000***** | **0.000***** | **0.001*** | 0.807 |
| **IL-10 (pg/mL)** | 98.22 ± 210.70 | 46.67 ± 28.60 | 19.58 ± 4.02 | **0.003**** | 0.964 | **0.002**** | **0.000***** |
| **MCP-1 (pg/mL)** | 207.28 ± 28.91 | 218.99 ± 50.22 | 236.05 ± 49.06 | 0.113 | 0.740 | 0.028 | 0.418 |
| **IGF-1 (pg/mL)** | 382.29 ± 378.92 | 156.86 ± 80.08 | 174.72 ± 218.77 | **0.000***** | **0.000***** | **0.000***** | 0.510 |
| **Humanin (pg/mL)** | 210.88 ± 53.30 | 188.16 ± 105.01 | 127..52 ±24.82 | **0.000***** | 0.775 | **0.000***** | 0.420 |
| **MOTSc (pg/mL)** | 572.82 ± 243.45 | 957.29 ± 416.01 | 909.17 ± 285.59 | **0.000***** | **0.000***** | **0.001**** | 0.807 |
| **p66Shc (pg/mL)** | 54.93 ± 17.26 | 50.88 ± 23.16 | 63.96 ± 15.99 | **0.002**** | **0.009**** | **0.012*** | **0.033*** |

**Abbreviations:** HbA1c – hemoglobin A1C, C5a – Complement component 5a, IL-6 – interleukin-6, TC – total cholesterol, HDL – high-density lipoprotein, LDL – high-density lipoprotein, 8-OHdG – deoxyguanosine, GSH – glutathione, GSSG – oxidized glutathione, IL-1β – interleukin-1β, IL-10 – interleukin-10, MCP-1 – monocyte chemoattractant protein-1, MOTSc – mitochondrial protein, p66Shc – adaptor protein.

*-p<0.05; **-p<0.01; ***-p<0.001

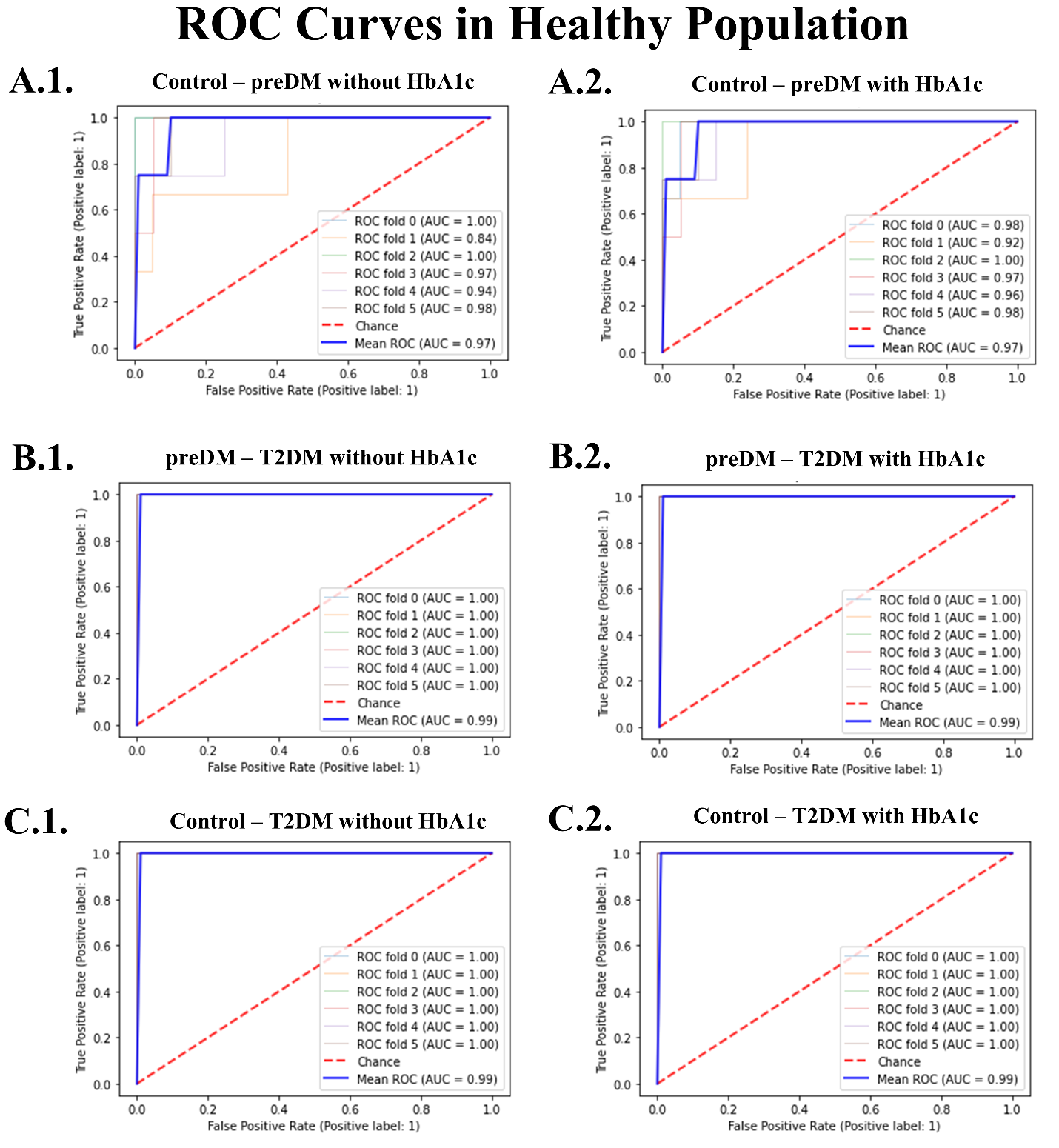

**Figure A1:** ROC curves in the healthy population with the use of LDA

b) Statistical analysis and classification ROC curves for the population with CVD and HT

**Table A2:** Characteristics and biomarker results of population with CVD and HT, in no diabetes, preDM, and T2DM subjects

|  |  |  |  | **Kruskal Wallis** | **Mann Whitney** | | |
| --- | --- | --- | --- | --- | --- | --- | --- |
|  | **No diabetes** | **preDM** | **T2DM** | **All Classes** | **No diabetes- preDM** | **No diabetes – T2DM** | **preDM – T2DM** |
|  | **Mean ± SD** | **Mean ± SD** | **Mean ± SD** | **p-value** | **p-value** | **p-value** | **p-value** |
| **N** | 95 | 26 | 95 |  |  |  |  |
| **Age (yr)** | 70.37 ± 8.84 | 69.96 ± 8.23 | 69.77 ± 8.45 | 0.764 | 0.654 | 0.488 | 0.990 |
| **HbA1c (%)** | 5.89 ± 0.32 | 5.88 ± 0.28 | 6.95 ± 0.95 | **0.000***** | 0.959 | **0.000***** | **0.000***** |
| **C5a (ng/mL)** | 6.74 ± 7.06 | 7.22 ± 4.32 | 12.68 ± 9.66 | **0.000***** | 0.379 | **0.000***** | **0.043*** |
| **D-Dimer (μg/L)** | 0.64 ± 0.29 | 0.61 ± 0.38 | 0.44 ± 0.37 | **0.000***** | 0.373 | **0.000***** | **0.015*** |
| **IL-6 (pg/mL)** | 26.57 ± 24.90 | 28.14 ± 36.76 | 31.55 ± 27.92 | 0.120 | 0.184 | 0.173 | 0.086 |
| **Triglyceride (mmol/L)** | 1.35 ± .52 | 1.65 ± 1.02 | 1.84 ± 0.91 | **0.001**** | 0.349 | **0.000***** | 0.159 |
| **TC (mmol/L)** | 4.98 ± 0.91 | 4.98 ± 0.91 | 4.09 ± 0.85 | **0.000***** | 0.842 | **0.000***** | **0.000***** |
| **HDL (mmol/L)** | 1.42 ± 0.32 | 1.30 ± 0.26 | 1.21 ± 0.31 | **0.000***** | 0.088 | **0.000***** | **0.034*** |
| **LDL (mmol/L)** | 2.95 ± 0.86 | 3.05 ± 0.90 | 2.09 ± 0.72 | **0.000***** | 0.513 | **0.000***** | **0.000***** |
| **8-isoprostane (mg/mmol)** | 0.99 ± 0.79 | 1.21 ± 0.93 | 4.26 ± 7.06 | **0.000***** | 0.103 | **0.000***** | **0.000***** |
| **8-OHdG (mg/mmol)** | 128.67 ± 93.88 | 107.83 ± 50.69 | 158.80 ± 137.62 | 0.118 | 0.649 | 0.086 | 0.095 |
| **GSH (mmol/L)** | 1603.89 ± 447.32 | 1781.27 ±390.75 | 1742.53 ±442.35 | 0.085 | 0.056 | 0.081 | 0.563 |
| **GSSG (mmol/L)** | 284.58 ± 135.89 | 282.33 ± 118.62 | 391.94 ± 273.03 | **0.002**** | 0.707 | **0.001**** | **0.032*** |
| **GSH/GSSG (-)** | 7.13 ± 4.18 | 7.64 ± 3.68 | 5.56 ± 2.22 | **0.019*** | 0.421 | **0.040*** | **0.008**** |
| **IL-1β (pg/mL)** | 6.03 ±5.32 | 5.62 ± 5.57 | 8.24 ± 11.64 | 0.486 | 0.405 | 0.609 | 0.229 |
| **IL-10 (pg/mL)** | 23.02 ± 23.53 | 18.45 ± 8.53 | 45.56 ± 62.01 | **0.022*** | 0.774 | **0.009**** | 0.097 |
| **MCP-1 (pg/mL)** | 251.18 ± 86.48 | 231.91 ± 6.27 | 207.81 ± 76.19 | **0.011*** | 0.275 | **0.003**** | 0.210 |
| **IGF-1 (pg/mL)** | 295.02 ± 134.07 | 345.82 ± 309.08 | 374.99 ± 422.20 | 0.243 | 0.847 | 0.099 | 0.396 |
| **Humanin (pg/mL)** | 241.91 ± 84.19 | 192.64 ± 66.40 | 298.26 ± 205.09 | **0.002**** | **0.003**** | 0.137 | **0.001**** |
| **MOTSc (pg/mL)** | 496.38 ± 150.66 | 613.70 ± 234.57 | 656.23 ± 263.59 | **0.000***** | **0.015**** | **0.000***** | 0.329 |
| **p66Shc (pg/mL)** | 44.69 ± 8.63 | 44.91 ± 9.53 | 45.25 ± 10.07 | 0.767 | 0.912 | 0.453 | 0.813 |

**Abbreviations:** HbA1c – hemoglobin A1C, C5a – Complement component 5a, IL-6 – interleukin-6, TC – total cholesterol, HDL – high-density lipoprotein, LDL – high-density lipoprotein, 8-OHdG – deoxyguanosine, GSH – glutathione, GSSG – oxidized glutathione, IL-1β – interleukin-1β, IL-10 – interleukin-10, MCP-1 – monocyte chemoattractant protein-1, MOTSc – mitochondrial protein, p66Shc – adaptor protein.

*-p<0.05; **-p<0.01; ***-p<0.001

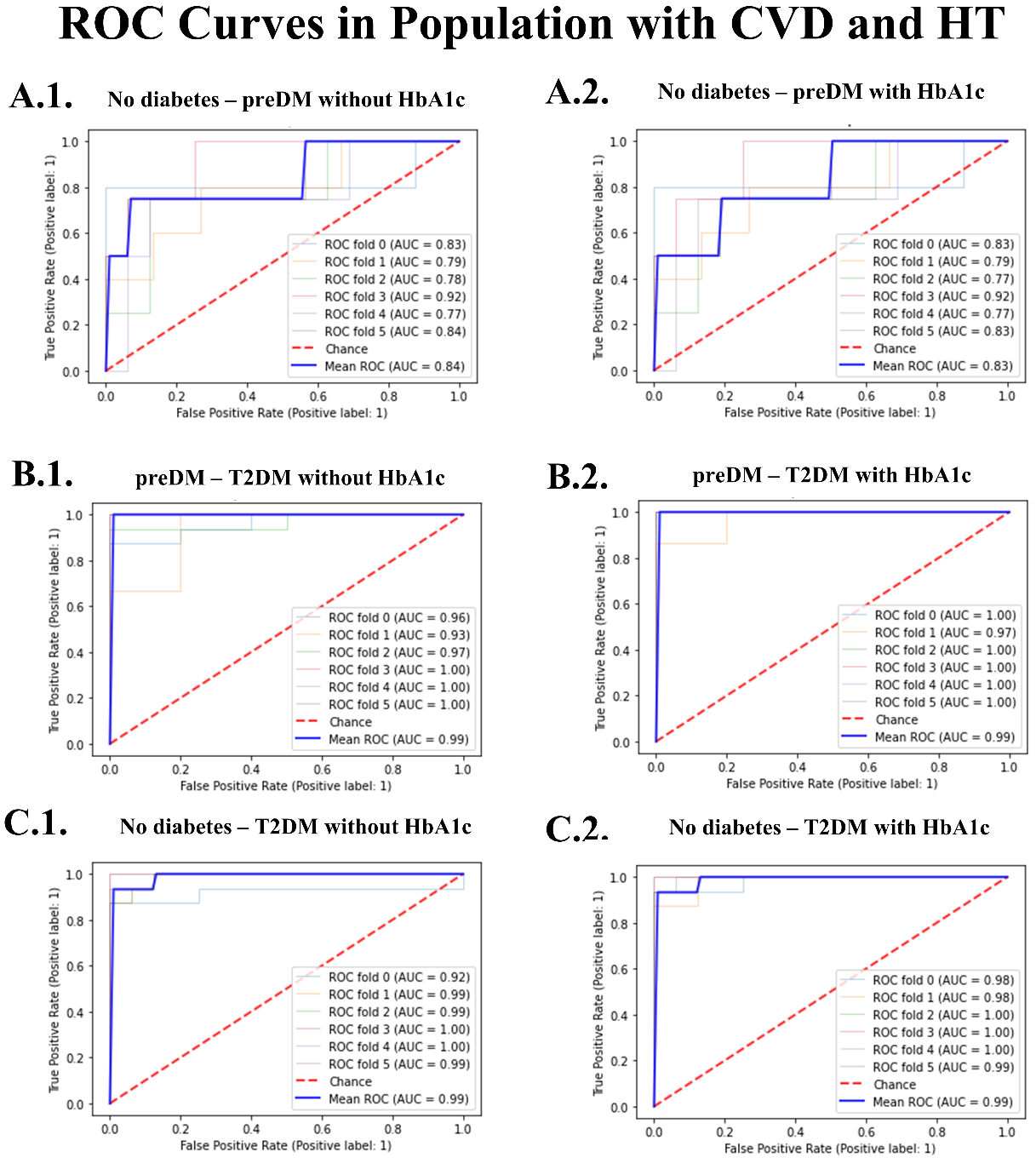

**Figure A2:** ROC curves in the HT and CVD population with the use of LDA.

c) Statistical analysis and classification ROC curves for the population with HT only

**Table A3:** Characteristics and biomarker results of population with HT only, in no diabetes, preDM, and T2DM subjects

|  |  |  |  | **Kruskal Wallis** | **Mann Whitney** | | |
| --- | --- | --- | --- | --- | --- | --- | --- |
|  | **No diabetes** | **preDM** | **T2DM** | **All Classes** | **No diabetes - preDM** | **No diabetes – T2DM** | **preDM – T2DM** |
|  | **Mean ± SD** | **Mean ± SD** | **Mean ± SD** | **p-value** | **p-value** | **p-value** | **p-value** |
| **N** | 80 | 20 | 48 |  |  |  |  |
| **Age (yr)** | 65.88 ± 11.03 | 68.35 ± 9.86 | 63.81 ± 9.64 | 0.168 | 0.413 | 0.176 | 0.071 |
| **HbA1c (%)** | 5.75 ± 0.33 | 5.86 ± 0.24 | 7.36 ± 13.29 | **0.000***** | 0.093 | **0.000***** | **0.000***** |
| **C5a (ng/mL)** | 8.93 ± 5.16 | 8.76 ± 3.58 | 18.27 ± 13.29 | **0.001**** | 0.382 | **0.001**** | **0.024*** |
| **D-Dimer (μg/L)** | 0.42 ± 0.29 | 0.40 ± 0.15 | 0.36 ± 0.18 | 0.244 | 0.558 | 0.152 | 0.186 |
| **IL-6 (pg/mL)** | 8.65 ± 6.13 | 16.67 ± 23.34 | 14.29 ± 19.39 | 0.820 | 0.806 | 0.537 | 0.752 |
| **Triglyceride (mmol/L)** | 1.50 ± 1.31 | 1.60 ± 1.26 | 1.89 ± 0.88 | **0.000***** | 0.261 | **0.000***** | **0.019*** |
| **TC (mmol/L)** | 5.28 ± 0.95 | 5.00 ± 0.75 | 4.70 ± 1.17 | **0.008**** | 0.220 | **0.003**** | 0.165 |
| **HDL (mmol/L)** | 1.40 ± 0.40 | 1.27 ± 0.31 | 1.36 ± 0.39 | 0.361 | 0.142 | 0.638 | 0.364 |
| **LDL (mmol/L)** | 3.32 ± 0.82 | 3.12 ± 0.71 | 2.53 ± 1.04 | **0.000***** | 0.300 | **0.000***** | **0.016**** |
| **8-isoprostane (mg/mmol)** | 2.31 ± 2.32 | 2.79 ± 2.14 | 2.23 ± 3.74 | **0.036*** | 0.181 | 0.064 | **0.022**** |
| **8-OHdG (mg/mmol)** | 178.16 ± 86.87 | 214.87 ± 106.73 | 87.72 ± 90.39 | **0.000***** | 0.140 | **0.000***** | **0.000***** |
| **GSH (mmol/L)** | 1759.39 ± 450.75 | 1703.88 ± 328.68 | 1476.18 ± 271.91 | **0.000***** | 0.688 | **0.000***** | **0.002**** |
| **GSSG (mmol/L)** | 335.36 ± 85.27 | 308.16 ± 81.66 | 232.46 ± 74.00 | **0.000***** | 0.096 | **0.000***** | **0.000***** |
| **GSH/GSSG (-)** | 5.53 ± 1.83 | 5.96 ± 1.96 | 6.84 ± 2.08 | **0.007**** | 0.326 | **0.002**** | 0.123 |
| **IL-1β (pg/mL)** | 11.82 ± 14.70 | 5.15 ± 4.88 | 7.48 ± 9.88 | **0.009**** | **0.012*** | **0.017*** | 0.548 |
| **IL-10 (pg/mL)** | 74.50 ± 120.50 | 17.24 ± 6.95 | 48.91 ± 124.83 | **0.000***** | **0.000***** | **0.001**** | 0.866 |
| **MCP-1 (pg/mL)** | 235.49 ± 82.61 | 184.32 ± 51.43 | 231.10 ± 43.66 | **0.016*** | **0.014*** | 0.492 | **0.005**** |
| **IGF-1 (pg/mL)** | 233.51 ± 157.78 | 314.64 ± 283.93 | 119.22 ± 101.64 | **0.000***** | 0.529 | **0.000***** | **0.001**** |
| **Humanin (pg/mL)** | 179.79 ± 67.12 | 147.03 ± 78.29 | 195.99 ± 55.38 | **0.015*** | **0.034*** | 0.092 | **0.011*** |
| **MOTSc (pg/mL)** | 649.02 ± 23.02 | 801.25 ± 234.70 | 887.85 ± 361.51 | **0.001**** | **0.018*** | **0.001**** | 1.000 |
| **p66Shc (pg/mL)** | 51.83 ± 12.74 | 51.12 ± 11.10 | 50.85 ± 11.22 | 0.905 | 0.959 | 0.739 | 0.596 |

**Abbreviations:** HbA1c – hemoglobin A1C, C5a – Complement component 5a, IL-6 – interleukin-6, TC – total cholesterol, HDL – high-density lipoprotein, LDL – high-density lipoprotein, 8-OHdG – deoxyguanosine, GSH – glutathione, GSSG – oxidized glutathione, IL-1β – interleukin-1β, IL-10 – interleukin-10, MCP-1 – monocyte chemoattractant protein-1, MOTSc – mitochondrial protein, p66Shc – adaptor protein.

*-p<0.05; **-p<0.01; ***-p<0.001

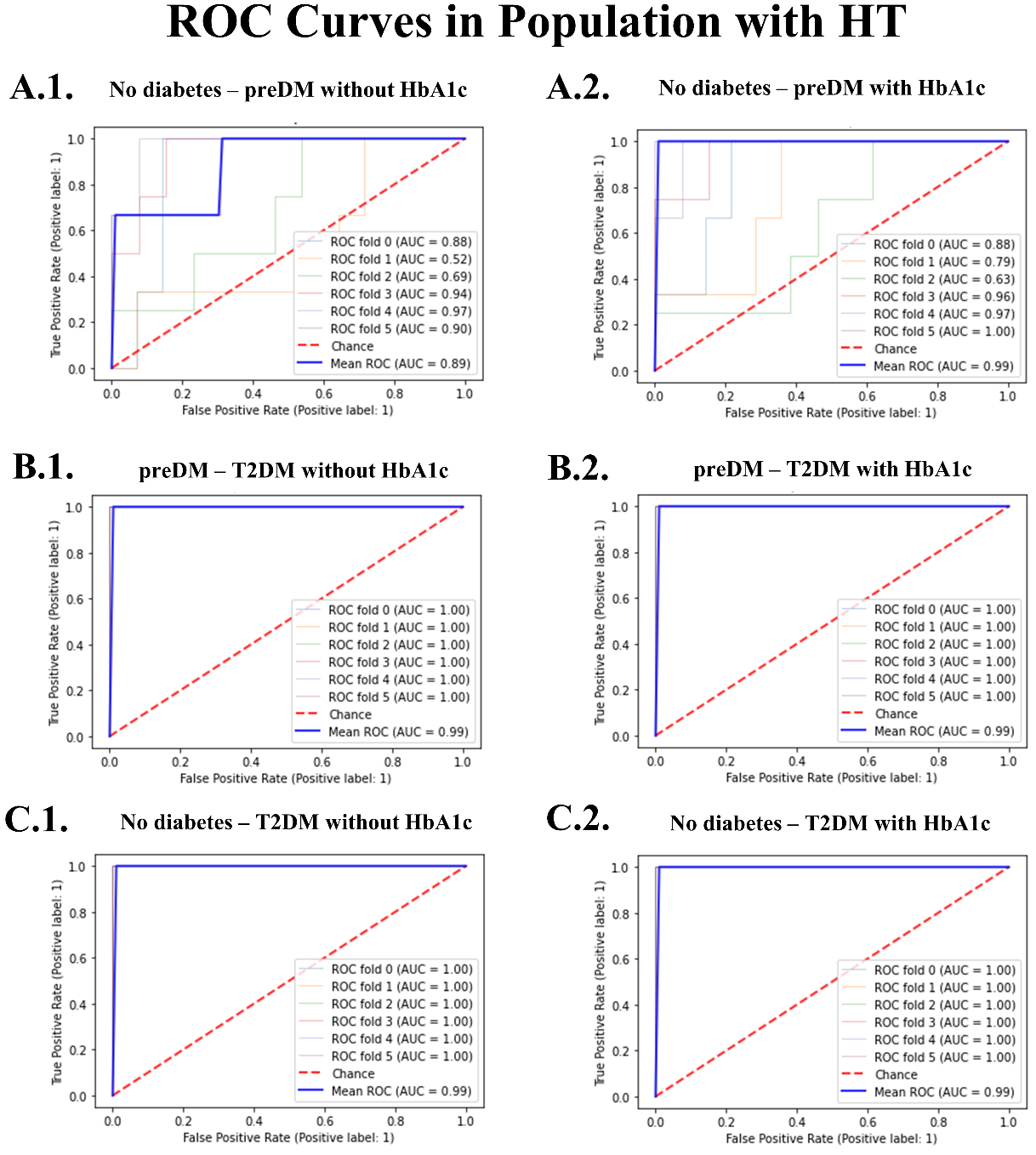

**Figure A3:** ROC curves in the HT population with the use of LDA.

d) Statistical analysis and classification ROC curves for the population with CVD only

**Table A4:** Characteristics and biomarker results of population with CVD only, in no diabetes, preDM, and T2DM subjects

|  |  |  |  | **Kruskal Wallis** | **Mann Whitney** | | |
| --- | --- | --- | --- | --- | --- | --- | --- |
|  | **No diabetes** | **preDM** | **T2DM** | **All Classes** | **No diabetes - preDM** | **No diabetes – T2DM** | **preDM – T2DM** |
|  | **Mean ± SD** | **Mean ± SD** | **Mean ± SD** | **p-value** | **p-value** | **p-value** | **p-value** |
| **N** | 34 | 13 | 14 |  |  |  |  |
| **Age (yr)** | 68.11 ± 11.14 | 67.69 ± 11.16 | 65.07 ± 10.31 | 0.607 | 0.821 | 0.296 | 0.680 |
| **HbA1c (%)** | 5.75 ± 0.23 | 5.79 ± 0.22 | 7.12 ± 0.64 | **0.000***** | 0.373 | **0.000***** | **0.000***** |
| **C5a (ng/mL)** | 8.16 ± 5.94 | 8.83 ± 8.66 | 8.44 ± 2.83 | 0.292 | 0.766 | 0.134 | 0.251 |
| **D-Dimer (μg/L)** | 0.37 ± 0.14 | 0.4 ± 0.14 | 0.40 ± 0.14 | 0.653 | 0.458 | 0.481 | 0.808 |
| **IL-6 (pg/mL)** | 14.11 ± 14.57 | 17.84 ± 15.02 | 6.42 ± 2.81 | **0.006**** | 0.422 | **0.004**** | **0.011*** |
| **Triglyceride (mmol/L)** | 1.56 ± 1.01 | 1.33 ± 0.77 | 1.72 ± 0.38 | 0.163 | 0.802 | 0.133 | 0.045 |
| **TC (mmol/L)** | 5.53 ± 1.01 | 5.08 ± 0.83 | 4.21 ± 0.91 | **0.001**** | 0.263 | **0.000***** | **0.021*** |
| **HDL (mmol/L)** | 1.42 ± 0.38 | 1.32 ± 0.25 | 0.99 ± 0.10 | **0.000***** | 0.493 | **0.000***** | **0.001**** |
| **LDL (mmol/L)** | 3.35 ± 0.75 | 3.13 ± 0.77 | 2.41 ± 1.88 | **0.007**** | 0.391 | **0.002**** | **0.039*** |
| **8-isoprostane (mg/mmol)** | 1.88 ± 0.93 | 1.98 ± 1.06 | 2.96 ± 1.88 | 0.316 | 0.857 | 0.124 | 0.366 |
| **8-OHdG (mg/mmol)** | 255.31 ± 216.68 | 269.29 ± 217.06 | 94.47 ± 31.47 | **0.016*** | 0.848 | **0.005**** | **0.043*** |
| **GSH (mmol/L)** | 1606.51 ± 555.98 | 1527.24 ± 594.49 | 2281.31 ± 455.42 | **0.002**** | 0.384 | **0.001**** | **0.007**** |
| **GSSG (mmol/L)** | 270.40 ± 87.71 | 309.86 ± 135.97 | 412.31 ± 79.48 | **0.000***** | 0.467 | **0.000***** | **0.028**** |
| **GSH/GSSG (-)** | 7.20 ± 5.51 | 6.98 ± 6.40 | 5.59 ± 0.72 | 0.357 | 0.346 | 0.345 | 0.230 |
| **IL-1β (pg/mL)** | 15.53 ± 13.76 | 12.72 ± 14.29 | 19.43 ± 15.13 | 0.146 | 0.252 | 0.224 | 0.060 |
| **IL-10 (pg/mL)** | 56.42 ± 49.39 | 59.80 ± 49.11 | 30.17 ± 42.63 | **0.002**** | 0.943 | **0.001**** | **0.005**** |
| **MCP-1 (pg/mL)** | 201.08 ± 51.89 | 210.79 ± 57.81 | 192.80 ± 79.82 | 0.784 | 0.784 | 0.609 | 0.511 |
| **IGF-1 (pg/mL)** | 295.90 ± 211.81 | 359.71 ± 208.71 | 129.23 ± 23.29 | 0.135 | 0.347 | 0.119 | 0.081 |
| **Humanin (pg/mL)** | 216.24 ± 58.39 | 197.19 ± 44.89 | 182.77 ± 81.01 | 0.594 | 0.408 | 0.776 | 0.337 |
| **MOTSc (pg/mL)** | 545.07 ± 105.54 | 593.17 ± 226.30 | 519.34 ± 59.97 | 0.732 | 0.517 | 0.724 | 0.474 |
| **p66Shc (pg/mL)** | 53.13 ± 16.80 | 54.36 ± 17.24 | 42.92 ± 6.05 | 0.634 | 0.932 | 0.443 | 0.306 |

**Abbreviations:** HbA1c – hemoglobin A1C, C5a – Complement component 5a, IL-6 – interleukin-6, TC – total cholesterol, HDL – high-density lipoprotein, LDL – high-density lipoprotein, 8-OHdG – deoxyguanosine, GSH – glutathione, GSSG – oxidized glutathione, IL-1β – interleukin-1β, IL-10 – interleukin-10, MCP-1 – monocyte chemoattractant protein-1, MOTSc – mitochondrial protein, p66Shc – adaptor protein.

*-p<0.05; **-p<0.01; ***-p<0.001

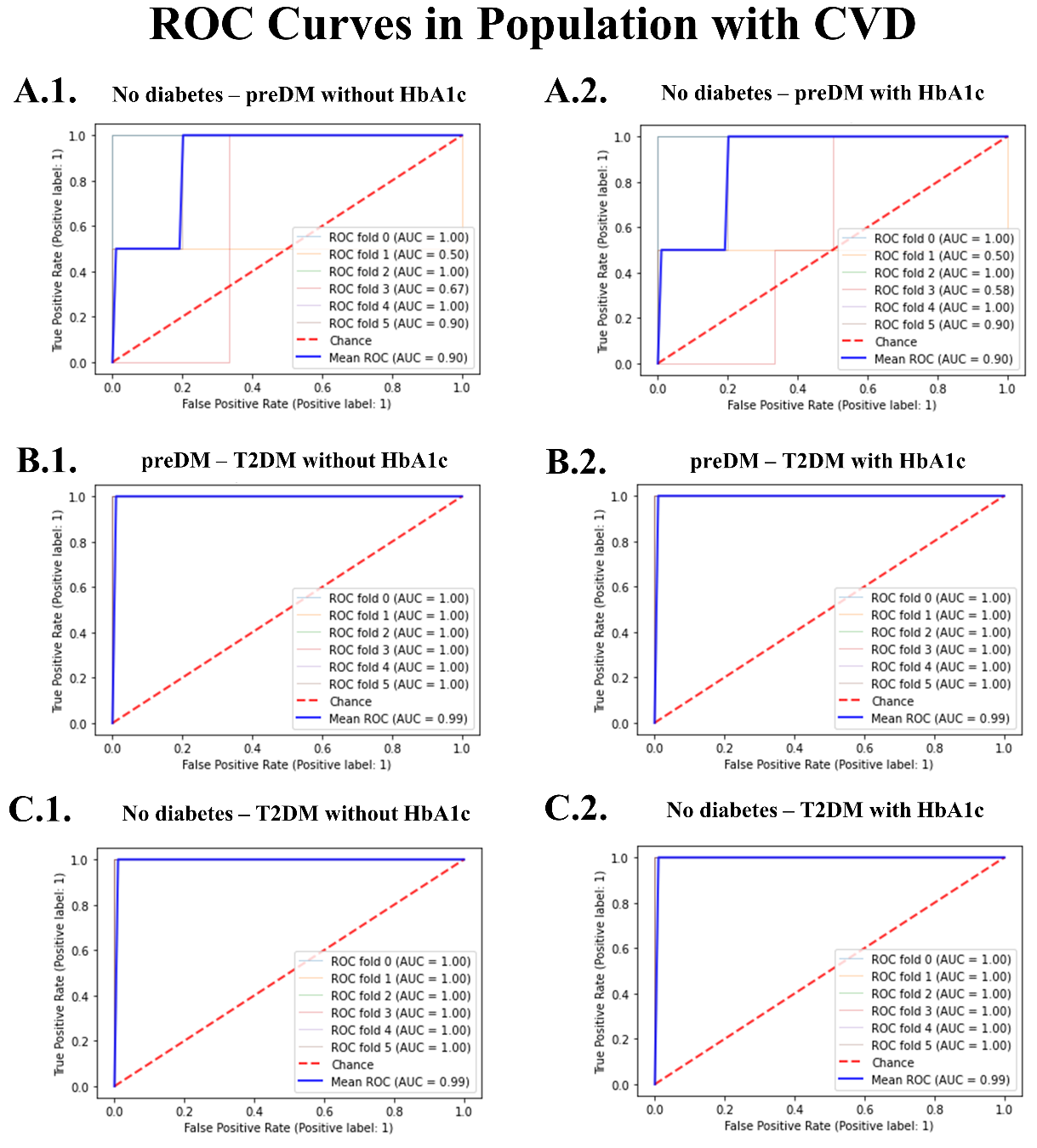

**Figure A4:** ROC curves in the CVD population with the use of LDA.
